# KLF6 and STAT3 co-occupy regulatory DNA and functionally synergize to promote axon growth in CNS neurons

**DOI:** 10.1101/257022

**Authors:** Zimei Wang, Vatsal Mehra, Matthew.T. Simpson, Brian Maunze, Lyndsey Holan, Erik Eastwood, Murray G. Blackmore, Ishwariya Venkatesh

## Abstract

Members of the KLF family of transcription factors can exert both positive and negative effects on axon regeneration in the central nervous system, but the underlying mechanisms are unclear. KLF6 and −7 share nearly identical DNA binding domains and stand out as the only known growth-promoting family members. Here we confirm that similar to KLF7, expression of KLF6 declines during postnatal cortical development and that forced re-expression of KLF6 in corticospinal tract neurons of adult female mice enhances axon regeneration after cervical spinal injury. Unlike KLF7, however, these effects were achieved with wildtype KLF6, as opposed constitutively active mutants, thus simplifying the interpretation of mechanistic studies. To clarify the molecular basis of growth promotion, RNA sequencing identified 454 genes whose expression changed upon forced KLF6 expression in cortical neurons. Network analysis of these genes revealed sub-networks of downregulated genes that were highly enriched for synaptic functions, and sub-networks of upregulated genes with functions relevant to axon extension including cytoskeleton remodeling, lipid synthesis and transport, and bioenergetics. The promoter regions of KLF6-sensitive genes showed enrichment for the binding sequence of STAT3, a previously identified regeneration-associated gene. Notably, co-expression of constitutively active STAT3 along with KLF6 in cortical neurons produced synergistic increases in neurite length. Finally, genome-wide ATAC-seq footprinting detected frequent co-binding by the two factors in pro-growth gene networks, indicating co-occupancy as an underlying mechanism for the observed synergy. These findings advance understanding of KLF-stimulated axon growth and indicate functional synergy of KLF6 transcriptional effects with those of STAT3.

**SIGNIFICANCE STATEMENT:** The failure of axon regeneration in the CNS limits recovery from damage and disease. These findings show the transcription factor KLF6 to be a potent promoter of axon growth after spinal injury, and more importantly clarify the underlying transcriptional changes. In addition, bioinformatics analysis predicted a functional interaction between KLF6 and a second transcription factor, STAT3, and genome-wide footprinting confirmed frequent co-occupancy. Co-expression of the two factors yielded synergistic elevation of neurite growth in primary neurons. These data point the way toward novel transcriptional interventions to promote CNS regeneration.

## INTRODUCTION

Promoting effective regenerative axon growth after central nervous system (CNS) injury remains a critical unmet goal in neuroscience research. The success of axon regeneration depends on both the extrinsic availability of growth-permissive tissue substrate (Geoffroy and Zheng, 2014; Silver et al., 2015), as well as the cell-intrinsic capacity to produce and assemble the diverse set of cellular materials necessary for axon extension (Blackmore, 2012; He and Jin, 2016; O’Donovan, 2016; Tedeschi and Bradke, 2017). Transcription factors (TFs), which can orchestrate broad patterns of gene expression, have long been recognized as key determinants of the success of axon regeneration (Smith and Skene, 1997; Blackmore, 2012; Venkatesh and Blackmore, 2017). It is increasingly clear that engaging multiple transcription factors and networks, as opposed to any single factor, will likely be necessary to fully restore regenerative capacity in adult neurons (Ma and Willis, 2015; Chandran et al., 2016; Venkatesh and Blackmore, 2017). The full set of TFs that regulate axon growth, and their optimal combinations, remain unclear.

The KLF family is comprised of 17 zinc finger transcription factors and play diverse roles in development, cellular differentiation, and stem cell biology (McConnell and Yang, 2010; Bialkowska et al., 2017). More recently, KLFs have emerged as important regulators of the neuron-intrinsic capacity for regeneration, with various family members displaying either anti-or pro-regenerative roles. For example, knockout of growth-inhibitory KLF4 promotes axon regeneration by retinal ganglion cells in the injured optic nerve (Moore et al., 2009; Qin et al., 2013). In contrast, forced expression of pro-regenerative KLF7 constructs improves axon regeneration by injured corticospinal tract (CST) neurons, and reduces dieback of axotomized sensory neurons (Blackmore et al., 2012; Wang et al., 2017). Importantly, exogenous KLF7 is poorly expressed in neurons, likely due to rapid protein degradation from an N-terminal targeting domain (Blackmore et al., 2012). Thus the pro-regenerative findings with KLF7 were obtained with a mutant form in which the N-terminal KLF7 domain was replaced with an artificial VP16 activation domain (VP16-KLF7). The molecular mechanisms that underlie growth regulation by KLF factors remain unclear.

Here we focus on KLF6, a close relative of KLF7, in order to clarify transcriptional mechanism that contribute to successful axon growth. We found that in contrast to previous findings with KLF7, wildtype KLF6 can be effectively overexpressed and produces a robust CST growth phenotype. We then used RNA-seq to identify changes in gene expression in KLF6-transduced cortical neurons, followed by network analysis to identify sets of genes relevant to axon extension. Next we used motif analysis of promoter sequences in KLF6-responsive genes, to identify candidate transcription factors that were predicted to co-regulate these genes. The recognition motif for STAT3, a known pro-regenerative transcription factor, was highly enriched in the promoter sequences of KLF6-responsive genes, suggesting potential functional interaction between these two factors. Consistent with this, we demonstrate synergistic gains in axon extension by forced co-expression of KLF6 and STAT3 in cultured neurons. Finally, using ATAC-sec footprinting analysis, we show high co-occupancy by KLF6 and STAT3 on regulatory DNA in regeneration-relevant gene networks, indicating co-binding as a likely mechanism for the observed synergy. These data demonstrate substantial growth promotion by KLF6 in CNS neurons, clarify the associated transcriptional changes, and demonstrate functional synergy with STAT3.

## METHODS

### Cloning and viral production

Mouse *KLF6* (accession BC020042) was purchased from Dharmacon and *caSTAT3* was a kind gift from Dr. Jessica Lerch (The Ohio State University) (Lerch et al., 2014). Genes were cloned into an AAV-compatible plasmid with CMV reporter (pAAV-MCS, Stratagene) using standard PCR amplification as in (Blackmore et al., 2012). AAV8-KLF6-2A-mCherry (red fluorescent protein) was constructed by excising VP16-KLF7 from the previously described VP16-KLF7-2A-mCherry vector (Blackmore et al. 2012) and replacing it with PCR-amplified KLF6. To create VP16-caSTAT3 the VP16 activation domain was PCR amplified from the pYSTrp3 vector (Invitrogen) and inserted N-terminal to caSTAT3. AAV production of AAV8-KLF6, and the previously described AAV8-EBFP and AAV8-EGFP (Blackmore et al., 2012), was performed at the Miami Project Viral Vector Core. All titers were brought to 5x10^13^ per ml.

### Spinal injury and retrograde identification of corticospinal neurons

All animal procedures were approved by the Marquette University Institutional Animal Care and Use Committee and complied with the National Institutes of Health Guide for the Care and Use of Laboratory Animals. Cervical dorsal hemisections were performed as in (Blackmore et al., 2012; Wang et al., 2015). Briefly, adult female C57/Bl6 mice (>8wks age, 20-22g) were anesthetized by Ketamine/Xylazine and the cervical spinal column exposed by incision of the skin and blunt dissection of muscles. Mice were mounted in a custom spine stabilizer. Using a Vibraknife device (Zhang et al., 2004), in which a rapidly vibrating blade is controlled via a micromanipulator, a transection was made between the 4^th^ and 5^th^ cervical vertebrae, extending from the midline beyond the right lateral edge of the spinal cord, to a depth of 0.85mm. To retrogradely identify CST neurons, immediately after injury a 2% solution of CTB-488 in sterile 0.9% NaCl (C22841-Thermofisher, Waltham, MA) was injected through a pulled glass micropipette fitted to a 10μl Hamilton syringe driven by a Stoelting QSI pump (catalog # 53311) and guided by a micromanipulator (pumping rate:0.04ul/min). 0.5μl of CTB-488 was injected at 2 sites located 0.5mm rostral to the injury, 0.2mm lateral to the midline, and to a depth of 0.8mm. Unilateral pyramidotomy was performed as described in (Blackmore et al. 2012, Wang et al. 2015). Briefly, a ventral midline incision was made to expose the occipital bone, the ventrocaudal part of which was removed using fine rongeurs. The dura was punctured and the right pyramid cut completely using a micro feather scalpel.

### Viral delivery to cortical neurons

Cortical neurons were transduced using intracerebral microinjection as described in (Blackmore et al., 2012; Wang et al., 2015). Mice were anesthetized with ketamine/xylazine (100/10 mg/kg, IP), mounted in a stereotactic frame, and skull exposed and scraped away with a scalpel blade. 0.5 μl of virus particles were delivered at two sites located 0 and 0.5mm anterior, −1.3mm lateral from Bregma and at a depth of 0.55mm, at a rate of 0.05 ul/min using a pulled glass micropipette connected to a 10µl Hamilton syringe driven by a programmable pump (Stoelting QSI). After each injection, the needle was left in place for 1 minute to minimize viral solution flow upwards.

### Horizontal ladder

Animals were pre-trained in two sessions on a horizontal ladder (30cm long, 1cm rung spacing) which included four crossing events across the ladder, until their performance plateued as previously described (Wang et al., 2015). Following gene treatment, the animals were tested 4, 8 and 12 weeks post treatment in four-crossing sessions with percent error quantified on the last three runs. All trials were video-recorded and errors were analyzed by blinded observers. Errors were defined as slips by the right forelimb (wrist breaks plane of ladder) and quantified as percentage of total steps by the right forelimb.

### Immunohistochemistry

For KLF6 expression profiling across development, brain tissue was snap-frozen in liquid nitrogen and cryosectioned (25 µm) and post-fixed for 20 mins with 4% Paraformaldehyde (PFA) in 1X PBS. Sections were incubated overnight with primary antibodies against KLF6 (SC R-173, Santa Cruz, Dallas, TX, 1:500 RRID: AB_2130443). For spinal injury experiments, adult animals were perfused with 4% paraformaldehyde in 1X-PBS (15710-Electron Microscopy Sciences, Hatfield, PA), brains and spinal cords removed, and post-fixed overnight in 4% PFA. Transverse brain sections and sagittal (spinal injury experiments) or transverse (pyramidotomy) sections of spinal cord were embedded in 12% gelatin in 1X-PBS (G2500-Sigma Aldrich, St.Louis,MO) and cut via Vibratome to yield 80µm sections. Sections were incubated overnight with primary antibodies PKCγ (SC C-19, Santa Cruz, Dallas, TX, 1:500, RRID:AB_632234) or GFAP (Z0334-DAKO, California, 1:500 RRID:AB_10013482), rinsed and then incubated for two hours with appropriate Alexafluor-conjugated secondary antibodies (Thermofisher, Waltham,MA, 1:500.) Fluorescent Images were acquired by an Olympus IX81 or Zeiss 880LSM microscopes.

### Quantification of axon growth

In the spinal cord injury experiment, axon growth was quantified from four 100µm sagittal sections of the spinal cord of each animal, starting at the central canal and extending into the right (injured) spinal tissue. The fiber index was calculated by dividing the spinal cord count by the total number of EGFP+ axons quantified in transverse sections of medullary pyramid, as in (Blackmore et al., 2012; Wang et al., 2015). The number of EGFP+ profiles that intersected virtual lines at set distances from the injury site or midline of the spinal cord, normalized to total EGFP+ CST axons in the medullary pyramid, was quantified by a blinded observer on an Olympus IX81 microscope (Liu et al., 2010; Blackmore et al., 2012; Geoffroy et al., 2015). Exclusion criteria for pyramidotomy experiments was animals with less than 80% decrease in PKCΓ in the affected CST, and for spinal injuries were 1) lesion depth less than 800μm 2) axons with straight white matter trajectory distal to the lesion, and 3) EGFP+ axons in thoracic cord (too far for regenerative growth).

### Western Blotting

Frontal cortex was dissected from early postnatal and adult mice and sonicated ~5s in 1ml RIPA lysis buffer (N653-Amresco,Solon,OH) with serine and cysteine protease inhibitors (4693159001-Roche, Boston, MA). After 30 minutes of incubation on ice, cell debris was removed by centrifugation at 21,000G at 4^°^ and protein concentration estimated using the BCA method. 25μg of protein was loaded and separated in a 4-20% SDS –PAGE gel (4561094-Bio-Rad, Hercules, CA) gel and transferred onto PVDF membrane (1620177-Bio-Rad, Hercules, CA) and blocked with 5% skim milk in 0.1% Tween 20 in PBS (TBST). The membranes were incubated with primary antibodies [mouse anti-KLF6 (SC-365633, Santa Cruz, Dallas, TX, 1:250)] in 5% skim milk in TBST overnight at 4^°^C. The membranes were incubated with HRP-conjugated secondary antibody (115035146-Jackson ImmunoResearch, West Grove, PA) at a 1:1000 dilution for 2 hours at room temperature. The labeled proteins were detected using the ECL agent (34077-Pierce, Rockford, IL), following the supplier’s manual. Briefly, a 1:1 solution of Supersignal west pico stable peroxide solution and enhancer solution was mixed and membranes were incubated in this solution for 7 minutes prior to detection. Detection and band quantification, was performed using an Odyssey Imaging system (LI-COR-Lincoln, NB) and band intensities were quantified using GAPDH (ab9484-Abcam, Boston, MA) as a reference with the built-in program. Quantification was the mean obtained from at least 3 biological replicates.

### Cortical cell culture and analysis of neurite outgrowth

All animal procedures were approved by the Marquette University Institutional Animal Care and Use Committee. Cortical neurons were prepared from a total of 15 litters of early postnatal (P3-P7) Sprague Dawley rat pups (Harlan), with each experimental replicate derived from a separate litter. A total of 15 litters were used across all experiments. Procedures for dissociation, transfection, immunohistochemistry, imaging, and neurite outgrowth analysis were performed as in (Simpson et al., 2015). Briefly, motor cortices were dissected, and neurons dissociated. Cells were transfected then plated at densities of 8,000 to 10,000 cells/well and maintained in culture for two to three days at 37 ^°^C, 5% CO2 in Enriched Neurobasal (ENB) media (10888-022-Thermofisher, Waltham, MA). All plates were pre-coated with poly-D-lysine hydrobromide (P7886-Sigma Aldrich, St.Louis, MO) followed by laminin (10µg/ml) (L2020-Sigma Aldrich, St. Louis, MO). Cultures were fixed with 4% PFA in 1X-PBS (15710-Electron Microscopy Sciences, Hatfield, PA), and immunohistochemistry was performed for βIII-tubulin (T2200-Sigma Aldrich, St. Louis, MO, 1:500) and/or KLF6 protein (sc-365633, Santa Cruz Biotechnology,Dallas,TX, 1:100) or NeuN (Millipore, MAB 377A5,1:250), with DAPI (2.5µg/ml in antibody solution) (D3571-Molecular Probes, Eugene,OR) to visualize nuclei. Automated microscopy acquired images and neurites were traced using Cellomics Scan v6.4.0 (Thermo Scientific,Waltham,MA). Average total neurite length was quantified for greater than 200 cells per treatment.

For the aggregated axon elongation assay, postnatal cortical neurons were first dissociated and transfected as described above. After transfection with test plasmids and an EGFP tracer, neurons were resuspended in Matrigel at a density of 133,000cells/ul, and two microliter aggregates were placed on in a 35mm petri dish, adjacent to a 2ul line of matrigel that contained no cells. Explants were covered in 3ml ENB media and maintained for 1 week, after which confocal microscopy produced an image that spanned the entire explant and extended 100um above the petri dish surface (11 optical sections, spaced 10um). The number of EGFP+ axons that crossed virtual lines at 1mm intervals from the edge of the cell aggregate was quantified.

### RNA-seq

P5 cortical neurons were prepared and maintained as described above. AAV8-EBFP or AAV8-KLF6 (titer-matched, 5*10^6^ total units) were added to each well at the time of plating (300,000 cells/well). To enrich for neurons, 100 nM 5-Fluoro-2’-deoxyuridine (FuDR) (F0503-Sigma-Aldrich, St.Louis,MO) was added 1 day post-transduction. NeuN immunohistochemistry confirmed >95% purity of nuclei (245 of 257 manually counted nuclei were NeuN+). At 3DIV, total RNA was extracted from neurons in culture using TRIzol reagent (10296028-Thermofisher, Waltham,MA) according to manufacturer’s instructions followed by DNAase-I treatment (EN052-Thermofisher, Waltham,MA). Total RNA quantification (Q32852-RNA HS assay, Qubit Fluorometer-ThermoFisher, Waltham,MA) and quality assessment (50671512-RNA nano chips, Bioanalyzer, Agilent, Santa Clara, CA) were performed according to guidelines recommended by ENCODE consortia (goo.gl/euW5t4). All RNA samples used for library construction had RIN scores >= 9. 100ng of total RNA from three separate cultures were used to construct replicate libraries for each treatment. The polyadenylated fraction of RNA was enriched by bead-based depletion of rRNA using TruSeq Stranded Total RNA Library Prep Kit with Ribo-Zero according to manufacturer’s instructions (RS-122-2201, Illumina technologies, San Diego,CA), and sequenced by Illumina HiSeq 2000 (University of Miami Genomics Core) to an average of 40 million paired-end reads. Preparation of cells for transduction and isolation of RNA were performed consecutively and by the same individuals to alleviate batch effects in sample preparation and sequencing.

Read quality post sequencing was confirmed using FASTQC package (per base sequence quality, per tile sequence quality, per sequence quality scores, K-mer content) (Brown et al., 2017). Trimmed reads were mapped to the rat reference genome [UCSC, Rat genome assembly: Rnor_6.0] using HISAT2 aligner software (unspliced mode along with–qc filter argument to remove low quality reads prior to transcript assembly) (Pertea et al., 2016). An average of 80-85% reads were present in map quality interval of >=30, and between 8-10% reads were excluded due to poor map quality (<10). Transcript assembly was performed using Stringtie and assessed through visualization on UCSC Genome Browser. Expression level estimation was reported as fragments per kilobase of transcript sequence per million mapped fragments (FPKM).

Differential gene expression analysis was performed using Ballgown software (Pertea et al., 2016). Isoforms were considered significant if they met the statistical requirement of having a corrected p-value of <0.05, FDR <0.05. Gene ontology analysis on the top 100 significant differentially expressed genes (up and down-regulated) was performed using default parameters in EnrichR software (Chen et al., 2013). Transcription factor binding site/motif analysis on KLF6 target gene promoters was performed using opossum v3.0 software (Kwon et al., 2012). Search parameters used were – JASPAR CORE profiles that scan all vertebrate profiles with a minimum specificity of 8 bits, conservation cut-off of 0.40, matrix score threshold of 90%, upstream/downstream of tss – 1000/300 bps, results sorted by Z-score >=10. Network analysis on significant differentially expressed genes (upregulated) was performed on the cytoscape platform v3.5.1 (Lotia et al., 2013), integrating multiple apps/plug-ins such as BiNGO, CyTransfinder, iRegulon, ClueGO, CluePedia and GeneMania (Maere et al., 2005; Bindea et al., 2009, 2013; Montojo et al., 2010; Lotia et al., 2013; Janky et al., 2014; Politano et al., 2016), using default parameters in a custom script.

### qPCR

To monitor developmental KLF6 expression, mouse motor cortices (P2, P8, P15, P21 and adult; 3 animals pooled per time-point) were homogenized in TRizol reagent (10296028-Thermofisher, Waltham,MA) and RNA was extracted via phase separation. The cDNA was synthesized from 500ng total RNA with Oligo dT priming using qSCRIPT reverse transcriptase (101414-112-Quanta Biosciences, Gaithersburg, MD). All quantitative qPCR experiments in this study were performed according to established MIQE guidelines(Bustin et al., 2009). All primer sequences and cycling conditions are available in **Sup Table 2**. A dissociation step was performed at the end of the amplification phase to confirm a single, specific melting temperature for primer pair. qPCR analysis was performed to determine relative levels of KLF6 transcript normalized to GAPDH internal reference using the following primer sequences – KLF6 5’ CACACAGGAGAAAAGCCTTACAGAT, 3’ GTCAGACCTGGAGAAACACCTG; GAPDH-5’ GCATCTTCTTGTGCAGTGCC, 3’ TACGGCCAAATCCGTTCACA. Relative gene expression was quantified using the 2^^^-(ΔΔCt) method (Livak and Schmittgen, 2001), ΔCt1 = Normalized Ct (KLF6 expression in P2) and ΔCt2 = Normalized Ct (KLF6 expression, time-points). Normalized gene expression data from 3-4 biological replicates were averaged and analyzed as fold change. For validation of RNAseq data, early postnatal (P3-P7) rat cortical neurons were dissociated, plated and maintained as described earlier. RNA extraction and cDNA synthesis was performed as described earlier. QPCR analysis was performed to determine the relative levels of four significant differentially expressed genes – s100a10, ccnd1, Lgals3, s100a11 mRNA, using GAPDH/beta-actin as an internal control. Primer sequences used are s100a11-5’ TCGAGTCCCTGATTGCTGTT, 3’ TCTGGTTCTTCGTGAAGGCG; ccnd1-5’ TCAAGTGTGACCCGGACTG, 3’ GGATCGATGTTCTGCTGGGC; s100a10-5’ CAGGGCCCAGGTTTCAACAG, 3’ GGAACTCCCTTTCCATGAGCA; Lgals3 - 5’ CCCAACGCAAACAGTATCAC, 3’ TTCTCATTGAAGCGGGGGTT. Relative gene expression was quantified as above, ΔCt1 = Normalized Ct (KLF6 over-expression) and ΔCt2 = Normalized Ct (EBFP overexpression). Normalized gene expression data from 3-4 biological replicates were averaged and analyzed as fold change.

### TF footprinting

Published ENCODE consortia ATAC-Seq datasets generated using postnatal day 0 mouse cortices were used for TF footprinting analyses (ENCSR310MLB) (Davis et al., 2018). Processed bam files were used as input for calling peaks using MACS2 software using the following parameters - macs2 callpeak -t <p0_atac>.bam -f BAM -g mm10 -n ATAC -q 0.01 (Zhang et al., 2008). Position Weight Matrices for KLF6 and STAT3 downloaded from Cis-BP database (Weirauch et al., 2014), narrow peak files generated using MACS2 and UCSC genome annotation mm10 were included as input to the CENTIPEDE footprinting algorithm, which was executed using default parameters and statistical thresholds (Pique-Regi et al., 2011). CruzdB package was used to annotate genomic loci containing footprints into either promoter (2000 bp upstream of TSS/300 bp downstream of TSS) regions or enhancer regions (intersect chromosomal locations with mouse ENCODE cis-element database) (Pedersen et al., 2013; Davis et al., 2018). Genes were separated into two categories –(1) Genes with only KLF6 footprints (2) Genes bound by both KLF6 and STAT3 in their promoter/enhancer regions, irrespective of binding distance. These genes were fed into the network analyses pipelines described above.

### Experimental Design and Statistical Analysis

For all cell culture experiments, neurite length from a minimum of 200 cells per treatment were averaged, and each experiment was repeated a minimum of three times on separate days. These averaged values were the basis for ANOVA with Sidak’s multiple comparisons. *In vivo* experiments were performed in a double-blind fashion, with non-involved lab personnel maintaining blinding keys. For *in vivo* experiments, animal numbers were determined by power analysis with G*Power software (α=.05, Power=0.8) using historical estimates of variability and adjusted for prior rates of animal attrition (Blackmore et al., 2012; Wang et al., 2015). Animals were randomized prior to viral treatment, with each surgical day including equal numbers from each group. The *in vivo* pyramidotomy experiment was initiated with ten female animals per group. One animal (control) was lost to mortality, and three animals (one control, two KLF6) animals were excluded on the basis of failed pyramidotomy (less than 80% reduction of PKCgamma signal in the affected tract). Spinal injury experiments were initiated with 18 control and 18 KLF6 animals. Three animals (two control, one KLF6) were lost to mortality, and three (two control, one KLF6) were excluded on the basis of insufficient injury depth and visibly spared axons. Axon counts (four sections per animal) at varying distances were compared between groups by RM ANOVA with Sidak’s multiple comparison’s test. For KLF6 expression across time, tissue was harvested from littermates at different time points. For the RNA-seq data validation, comparisons in fold change were analyzed with One-way ANOVA with Sidak’s multiple comparison test. For all software used for bioinformatic analyses, default inbuilt statistical models were employed as described above. All statistical tests were performed with Graphpad Prism software v7.0(San Diego, USA) and all error bars are ±SEM and statistical significance was accepted with p-value <0.05.

## RESULTS

### Cortical KLF6 expression declines developmentally and is not increased by spinal injury

Consistent with a positive role in axon growth, KLF6 transcript is highly expressed in the embryonic brain during periods of axon growth and then developmentally downregulated throughout the CNS during postnatal development, including in CST neurons (Laub et al., 2001; Arlotta et al., 2005). A more recent study, however, reported detection of KLF6 protein by immunohistochemistry in the adult brain (Jeong et al., 2009). We therefore re-examined KLF6 expression at the protein level across early postnatal development using immunohistochemistry western blotting, and quantitative PCR. As expected, KLF6 was readily detectable by immunohistochemistry (red) in early postnatal forebrain neurons (**Fig. 1A**), and western blot analysis of whole cortex detected protein of ~40kd, consistent with KLF6’s expected size (Kimmelman et al., 2004) (**Fig. 1A,C**). In the adult, KLF6 was also detectable by immunohistochemistry (red), but with intensity much dimmer than postnatal tissue processed identically (**Fig. 1B**). KLF6 was also detected by western blot in the adult cortex, with expression about 10% of that in postnatal cortex (**Fig. 1C**) (p<.001, ANOVA with post-hoc Tukey’s). Consistent with this, qPCR analysis of P1, P8, P15, P21, and adult cortex showed a 70% reduction in KLF6 transcript during early postnatal development (p<.01, ANOVA with post-hoc Tukey’s, **Fig. 1E**). Combined, these data confirm a substantial downregulation of KLF6 protein and transcript in the developing cortex.

**Figure 1.**
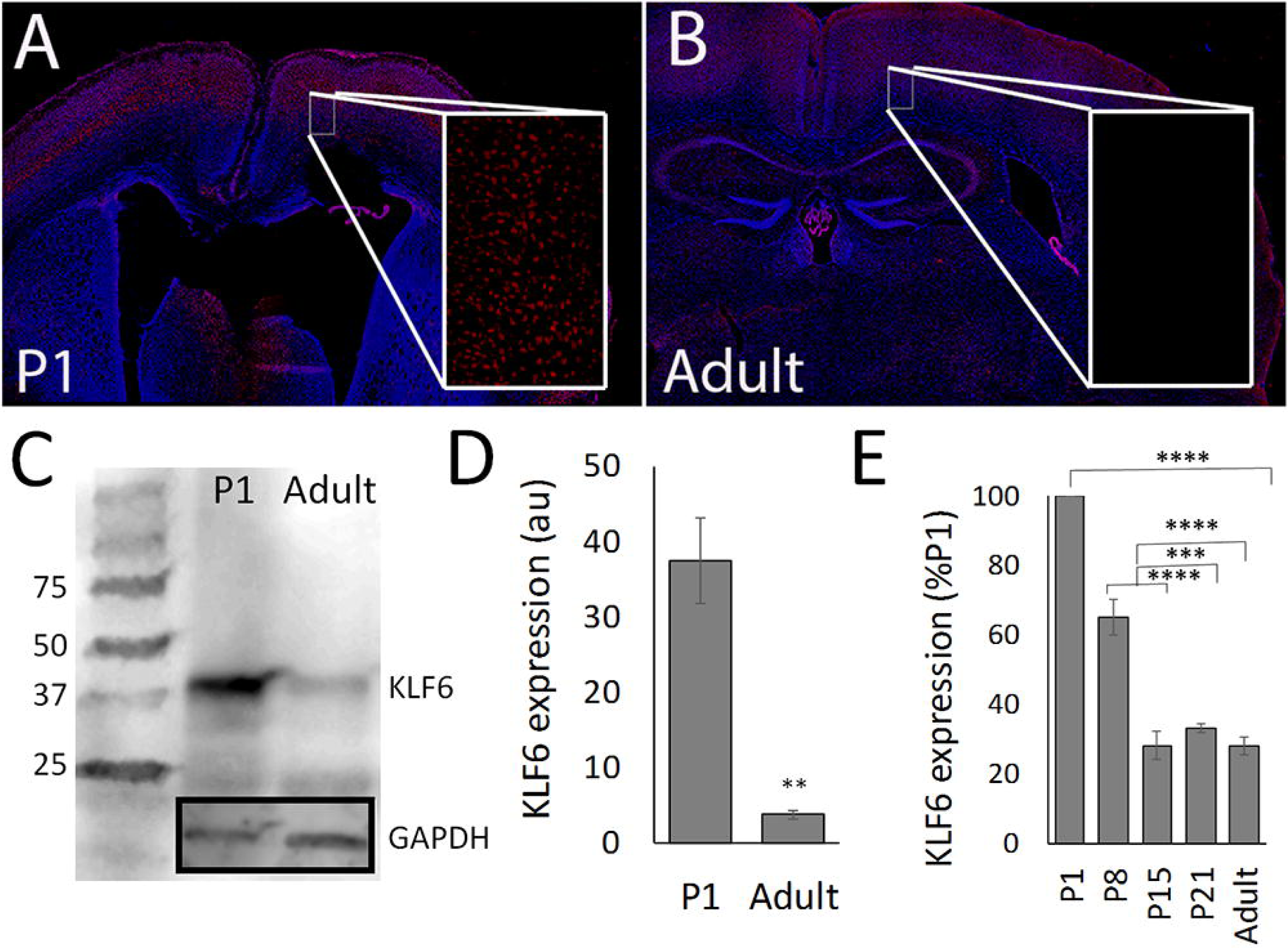
KLF6 expression in the cortex decreases during postnatal development. (A,B) Transverse sections of motor cortex were prepared and immunohistochemistry for KLF6 performed. Nuclear KLF6 signal (red) is visible at postnatal day 1 (P1, A), but dimmer in the adult (B). (C) Western blotting for KLF6 in whole cortex reveals bands of the expected size and a large reduction in KLF6 signal in adult tissue compared to P1. (D) Quantification of three replicate blots from three different sets of animals shows a significant reduction in KLF6 signal, normalized to GAPDH loading controls (**p<.05, ANOVA with Tukey, N=3). (E) Quantitative PCR analysis of whole cortex shows a significant reduction in KLF6 transcript abundance during postnatal development (***=p<.001 ANOVA with Tukey, N=3 per sample). Scale bars are 1mm in main panels, 50µm in insets.

To determine whether KLF6 expression in CST neurons is affected by spinal injury, fifteen animals received sham or cervical C4/5 hemisection and injections of the retrograde tracer CTB-488 to cervical spinal cord. Animals were sacrificed between three days to four weeks post-injury (3 animals per time point), and retrogradely identified CST neurons in the cortex were examined with immunohistochemistry for KLF6. Immunohistochemistry was performed simultaneously on all tissue sections, which were visualized with identical acquisition parameters. To account for residual variability, KLF6 intensity in the nuclei of identified CST neurons was normalized to the within-slice average intensity of non-injured cells in layer 4 (**Fig. S1-A**). KLF6 signal remained dim in injured CST neurons at all time-points post-injury (**Fig. S1 C-H**). Thus, similar to KLF7 and other regeneration-associated genes (Mason et al., 2003; Moore et al., 2009; Blackmore et al., 2012), KLF6 is developmentally downregulated and expression levels remain depressed in adult CST neurons after spinal injury.

### Forced KLF6 expression promotes CST sprouting and regeneration after spinal injury

We next tested the effects of overexpressing KLF6 on neurite outgrowth in postnatal rat cortical neurons, a well-characterized model system (Blackmore et al., 2010; Simpson et al., 2015). Rat neurons were used for cell culture experiments to improve per-animal cell yield and viability compared to mice; gene function in this rat system has repeatedly proven to predict pro-regenerative activity in mice (Moore et al., 2009; Blackmore, 2012; Blackmore et al., 2012). First, to establish baseline expression of KLF6 expression, cortical neurons were cultured for two days and immunohistochemistry was performed with antibodies to KLF6 and neuron-specific β-III tubulin. High content-screening microscopy was used to quantify KLF6 signal (**Fig. S2-E**) and neurite growth in thousands of individual β-III+ neurons (**Fig. S2-D**). Neurons displayed a range of KLF6 signal intensity. Consistent with a pro-growth role for KLF6, neurons in the highest quartile of KLF6 intensity produced neurites that were significantly longer than the lowest quartile (top quartile 116.2% of bottom, p<.001, ANOVA with post-hoc Sidak’s) (**Fig.S2-A,B**).

To test more directly the effect of KLF6 overexpression, postnatal cortical neurons were transfected with plasmid DNA encoding KLF6 or EBFP (enhanced blue fluorescent protein) control, and automated microscopy was used to quantify neurite outgrowth (**Fig.S2-F**). To distinguish transfected cells a fluorescent EGFP (enhanced green fluorescent protein) reporter was co-transfected at a ratio of 1 part EGFP to 4 parts test plasmid; this ratio was previously established to result in greater than 90% co-expression (Blackmore et al., 2010). To determine how the response to KLF6 might vary during early postnatal development, neurons were collected at either postnatal day 3 (P3) or at P7, the oldest age at which neurons could be reliably harvested and transfected in our hands. Notably, in baseline control conditions P3 neurons extended neurites that were significantly longer than P7 (**Fig.S2-H**). At P3, KLF6 overexpression produced a non-significant trend towards increased neurite length, but significantly increased neurite lengths in P7 neurons (**Fig.S2-H**) (EBFP 201.7 ± 17.5 µm, KLF6 283.4 ± 28.2 µm p<.05, ANOVA with Sidak’s). We attempted to simplify labeling of KLF6-expressing cells by creating a construct in which a T2A “self cleaving” oligopeptide linker was inserted between KLF6 and an mCherry red fluorescent protein (de Felipe et al., 2006). This approach creates two distinct proteins from a single vector and has been used by our lab and others to study regeneration-associated genes (Blackmore et al., 2012; Lerch et al., 2014; Wang et al., 2015). We found, however, that that growth promotion by KLF6-2A-mCherry was substantially lower than untagged KLF6 (KLF6 137% ± 2.7SEM, p<.01 versus mCherry control; KLF6-2A-mCherry 111% ± 7.9SEM, p>.05 versus mCherry control, ANOVA with post-hoc Sidak’s). The mechanism for this difference is unclear, but based on these data we elected to use untagged KLF6 constructs for all subsequent experiments, relying on EGFP from a separate plasmid to mark transfected neurons. Cell-based HCS analysis of immunohistochemistry confirmed increased expression of KLF6 in transfected neurons (**Fig.S2-G**). Collectively these data establish a correlation between endogenous KLF6 expression and total neurite length in cortical neurons, and confirm that elevated KLF6 expression is sufficient to enhance neurite length in postnatal cortical neurons.

We next tested the ability of forced KLF6 expression to enhance sprouting by adult CST neurons *in vivo*. Twenty adult female mice received unilateral pyramidotomy to deprive the right side of the spinal cord of CST input, followed by cortical injections of AAV8-EBFP control or AAV8-KLF6 to the right cortex in order to target uninjured CST neurons that project to the left spinal cord. AAV8-EGFP was co-injected as an axon tracer, using a 1:3 ratio shown previously to result in >90% co-expression (Blackmore et al., 2012; Wang et al., 2015). Twelve weeks later, transverse sections of cervical spinal cord were prepared, and PKCgamma immunohistochemistry was performed to confirm confirmed successful ablation of the left CST (**Fig. 2A,B insets**) (Liu et al., 2010; Blackmore et al., 2012; Geoffroy et al., 2015). One animal (control) was lost to mortality, and three animals (one control, two KLF6) were excluded for incomplete pyramidotomy. The total number of EGFP+ CST axons was similar between groups (EBFP: 2031.3 ± 158.6 SEM, KLF6: 2168.0 ± 161.3 SEM, p>0.05, t-test). Compared to EBFP control, KLF6 expression caused significant elevation of cross-midline sprouting by CST axons at distances up to 600µm from the midline (N=8 per group, p<.05 RM ANOVA with Sidak’s multiple comparison **Fig. 2C**). To test whether this enhanced sprouting was accompanied by improvements in forelimb function, animals were tested on a horizontal ladder task, which quantifies the percent of steps by the affected right forelimb that are mistargeted. As expected, pyramidotomy injury significantly increased the slip rate, but animals treated with KLF6 did not differ significantly from control animals at any timepoint post-injury (**Fig. 2D**). Thus forced KLF6 expression enhances compensatory sprouting by intact CST neurons, but does not yield improved forelimb targeting in the horizontal ladder task.

**Figure 2.**
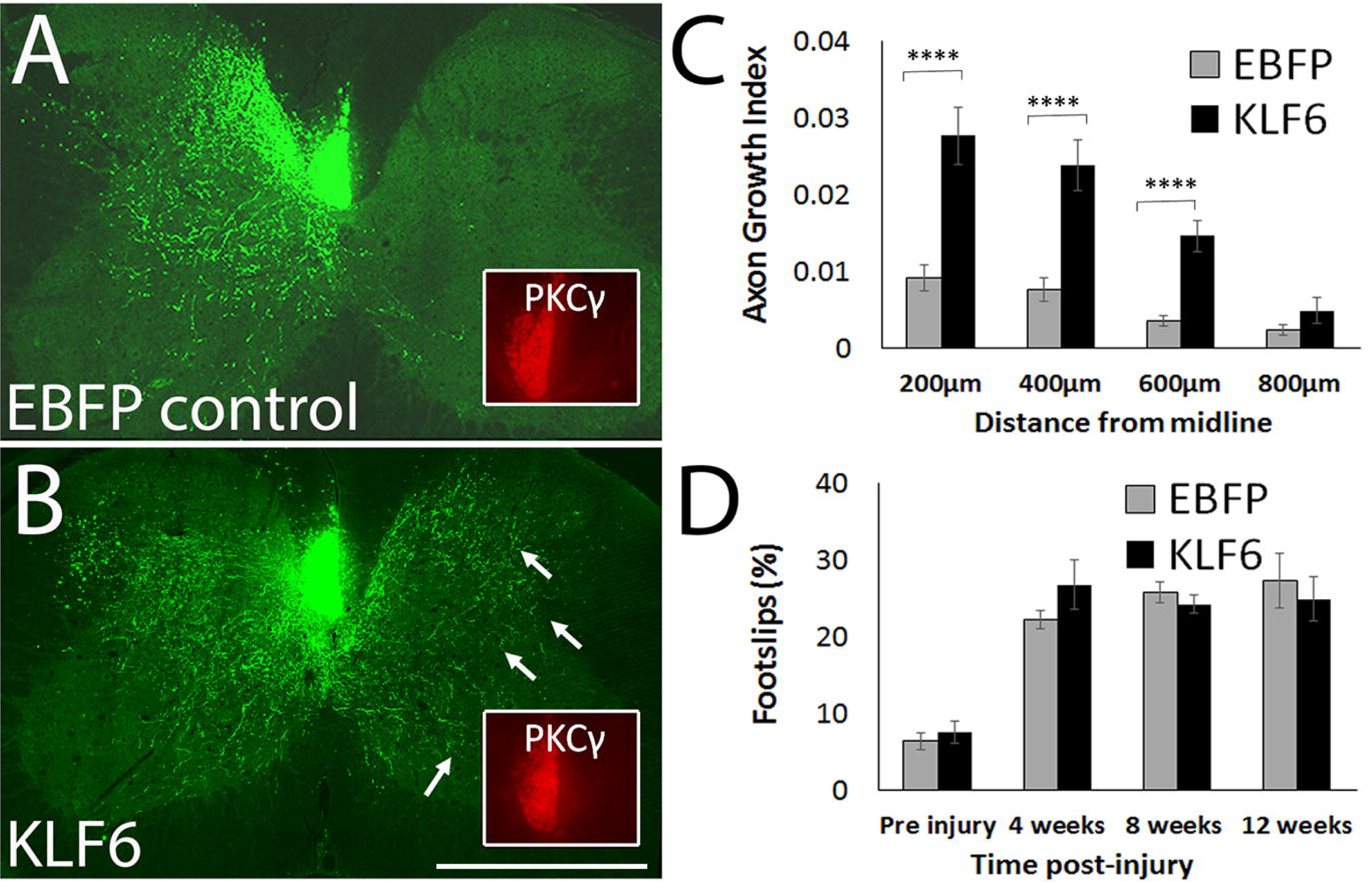
Forced KLF6 expression enhances CST cross-midline sprouting after unilateral pyramidotomy. Adult mice received unilateral pyramidotomy to deprive the right spinal cord of CST input, followed by cortical injection of AAV8-KLF6 or AAV8-EBFP control mixed with AAV8-EGFP tracer. (A,B) show transverse sections of cervical spinal cord twelve weeks post-injury. In EBFP-treated animals, left CST axons (green) show low levels of sprouting across the midline, whereas in KLF6-treated animals numerous CST axons are visible in the right spinal (arrows, B). Pyramidotomy was confirmed by PKCΓ immunohistochemistry (red, insets). (C) CST axon growth was quantified by counting the number of CST axons (EGFP+, green) that intersect virtual lines at set distances to the right of the midline, normalized to the total number of EGFP+ axons present in the medullary pyramids. Axon growth was significantly elevated in KLF6-treated animals at distances up to 600µm from the midline. (N=8 animals per group,**=p<.001, RM ANOVA with Sidak’s multiple comparison’s test). (D) Right forelimb function was tested by quantifying the percent foot-slips as animals traversed a horizontal ladder. Errors were increased by injury but unaffected by KLF6 treatment (p>.05, RM ANOVA). Scale bar is 1mm.

Next, to test the effect of forced KLF6 expression on regeneration by directly injured neurons, AAV8-EGFP tracer and AAV8-EBFP or AAV8-KLF6 were delivered to the left cortex of adult female mice (18 animals per group). One week later, CST axons in the right cervical spinal cord were transected by a dorsal hemisection injury (**Fig. 3A**). Animals were sacrificed twelve weeks later, and immunohistochemistry readily confirmed expression of KLF6 protein in sites of AAV injections, but not in AAV8-EBFP animals (**Fig. 3B,C**). Sagittal sections of cervical spinal cord were prepared, and GFAP immunohistochemistry (blue) was used to visualize the site of injury. Three animals (two control, one KLF6) were lost to mortality, and three (two control, one KLF6) were excluded on the basis of insufficient injury depth and visibly spared axons. To assess axon regeneration, the number of EGFP-labeled CST axons, normalized to the total number EGFP-labeled CST axons in the medullary pyramid, was quantified at set distances distal to the injury. The total number of EGFP+ CST axons detected in the medullary pyramids was similar between groups (EBFP: 2681.9 ± 155.4 SEM, KLF6: 3228.5 ± 240.4 SEM, p>0.05, t-test). As expected, EBFP control animals displayed very little CST growth distal to the injury site (**Fig. 3D**) In contrast, KLF6-treated animals showed significantly elevated CST growth up to 3mm beyond the injury site (N=14 EBFP control, 16 KLF6, p<.05 RM ANOVA with Sidak’s multiple comparison, **Fig. 3E,F**). Similar to the pyramidotomy experiment, however, KLF6 treatment had no effect on forelimb placement in the horizontal ladder task at any time point post-injury (**Fig. 3G**). Overall, these data indicate that forced re-expression of KLF6 in CST neurons promotes axon sprouting and regeneration, but is not sufficient to improve forelimb placement.

**Figure 3.**
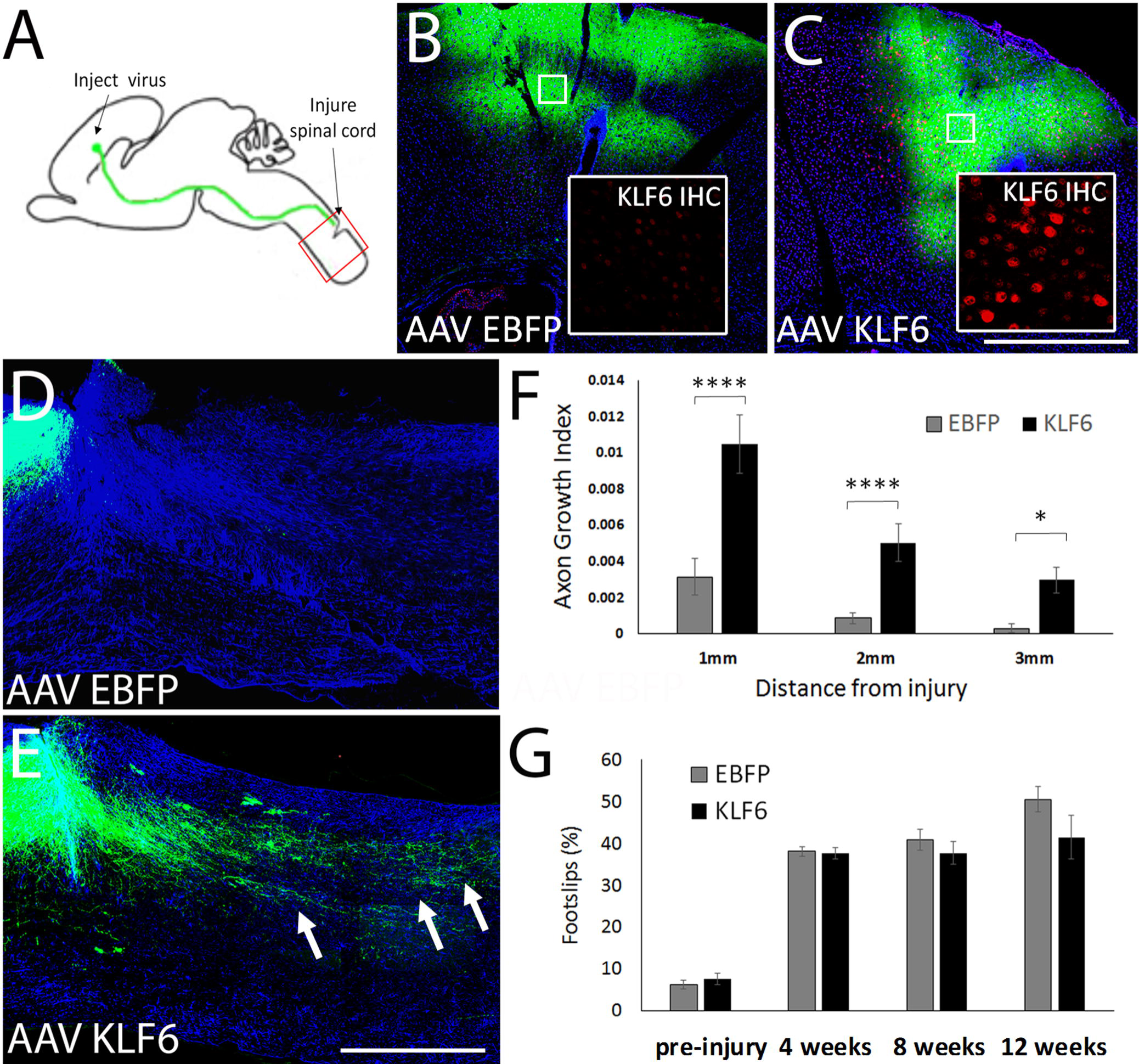
KLF6 overexpression increases CST axon growth after spinal injury. (A) Mice received cortical injection of AAV8-KLF6 or AAV8-EBFP control, mixed 2:1 with AAV8-EGFP tracer. One week later CST axons were severed by C4/5 dorsal hemisection. (B-C) Twelve weeks post-injury, KLF6 protein expression (red) was readily detectable in viral-injected regions of cortex in AAV8-KLF6 treated animals (C, inset) but not in AAV8-EBFP treated animals. (D,E) Sagittal sections of spinal cord were prepared and GFAP immunohistochemistry (blue) performed to define injury sites. EGFP-labeled CST axons (green) were rarely visible in tissue distal to the injury in AAV8-EBFP treated animals, but often extended for several millimeters in AAV8-KLF6 animals (arrows, E). (F) Quantification of the number of CST axons that intersect virtual lines below the injury, normalized to total EGFP+ axons in the medullary pyramid, showed significant elevation of axon growth in AAV8-KLF6 animals. N=14 control and 16 KLF6 animals, ****=p<.0001,*=P<0.05, RM 2-way ANOVA with Sidak’s multiple comparisons test. (G) Right forelimb function was tested by quantifying the percent foot-slips as animals traversed a horizontal ladder. Errors were increased by injury but unaffected by KLF6 treatment (p>.05, RM ANOVA). Scale bars are 1mm (B,C,D,E), 20µm in insets (B,C)

### RNA-seq analysis of transcriptional effects of KLF6 overexpression

Although KLF transcription factors have been functionally linked to regenerative axon growth, the underlying molecular mechanisms remain unclear. We therefore adopted an RNA-seq approach to examine the transcriptional consequences of forced KLF6 overexpression. Postnatal cortical neurons were treated with AAV8-mCherry control or AAV8-KLF6 virus, using titers that resulted in >95% efficiency of transduction. In addition, FUDR was added to remove contaminating glia, resulting in >95% neuronal purity as assessed by NeuN immunoreactivity. Three days later, RNA (RNA Quality Score RIN >9.0) was collected from three experimental replicates from three separate litters, and deep sequencing performed with Illumina HiSeq 2000, with an average of 40 million 100bp reads per sample. After extensive quality control in accordance with ENCODE guidelines (RNA integrity, library quality, sequence read quality and distribution) and alignment to the rat genome (see methods and **Fig. S3**), differential gene expression was assessed using Ballgown software. KLF6 overexpression significantly altered the expression of 454 genes (p-value<0.05, FDR<0.05), 55% of which were upregulated and 45% down-regulated. The genome-wide fpkm values for all samples and the complete set of differentially expressed transcripts are available in (**Sup_Table 1**).

To validate the RNA-seq data, newly prepared cortical neurons were transfected with KLF6 or EBFP control, and qPCR was used to assess changes in genes predicted by RNA-seq to be most affected by KLF6 expression. As an additional control, qPCR for putative KLF6-downstream genes was also performed after transfection with mutant forms of KLF6. Transcriptional activity of KLF6 and −7 has been previously shown to be sensitive to acetylation of two lysines (K) located in the first zinc finger of the DNA binding domain (Li et al., 2005; Wang et al., 2016). Accordingly, lysines 209 and 213 of KLF6 were mutated to either arginine (R) or glutamine (Q) to block or mimic the effects of acetylation, respectively (**Fig. S4-A**). In assays of neurite outgrowth, mutation to glutamine (KLF6-K-to-Q) completely blocked growth promotion by KLF6, whereas transfection with the arginine mutant (KLF6-K-to-R) elevated growth promotion modestly but significantly above the level of wildtype KLF6 (**Fig.S4-B**). qPCR analysis showed that transfection with either wildtype KLF6 or KLF6-K-to-R significantly upregulated *s100a10, ccnd1, Igals3*, and *s100a11*, with fold changes similar to those in the RNA-seq dataset (**Fig. S4-C**). In contrast, transfection with KLF6-K-to-Q showed no significant elevation in putative KLF6 target genes (**Fig.S4-C**). qPCR for KLF6 confirmed successful transfection with all constructs (fold increases of 189.2 ± 43.05 SEM, 631.28 ± 101 SEM, 434.42 ± 276 SEM, for wildtype, K-to-Q, and K-to-R mutants respectively). Overall these data are consistent with the previously established importance of these zinc finger lysines for KLF6 activity (Li et al., 2005), and serve to independently validate the RNA-seq-based identification of differentially expressed genes.

As a first step in clarifying the mechanisms by which KLF6 affects neurite outgrowth we performed gene network analysis of the genes most up- and down-regulated by KLF6. Network analysis identified 5 distinct subnetworks within the upregulated cohort. Interestingly, genes within each subnetwork were highly enriched for distinct functions associated with axon extension, including regulation of cytoskeleton remodeling, regulation of cellular component size, lipid synthesis and transport, membrane raft assembly, and bioenergetics (**Fig. 4A**). In contrast, we identified 3 distinct sub-networks within the downregulated gene cohort, all enriched for functions related to synaptic transmission (**Fig.4B**). Overall these data indicate that KLF6 expression may affect axon growth primarily by altering the abundance of actin regulatory genes, while also engaging complementary pro-growth mechanisms. To test whether forced expression of cytoskeletal-interacting genes might phenocopy KLF6 growth promotion, in follow-up experiments the cytoskeletal-interacting genes most affected by KLF6 were overexpressed in cortical neurons. Unlike KLF6, however, expression of these putative effector genes did not significantly alter neurite lengths (Anxa1 107.5±7.5%, Anxa2 100.5±2.7%, S100a4 97.4±10.0%, S100a10 95.8±3.5%, N>200 cells in three experiments, p>.05 ANOVA with post-hoc Sidak’s). One likely explanation is that growth promotion by KLF6 may require changes in the abundance of multiple, functionally interacting genes. If so, single or even multi-expression of putative target genes may not be a feasible strategy to mimic the KLF6 effect by engaging downstream effectors.

**Figure 4.**
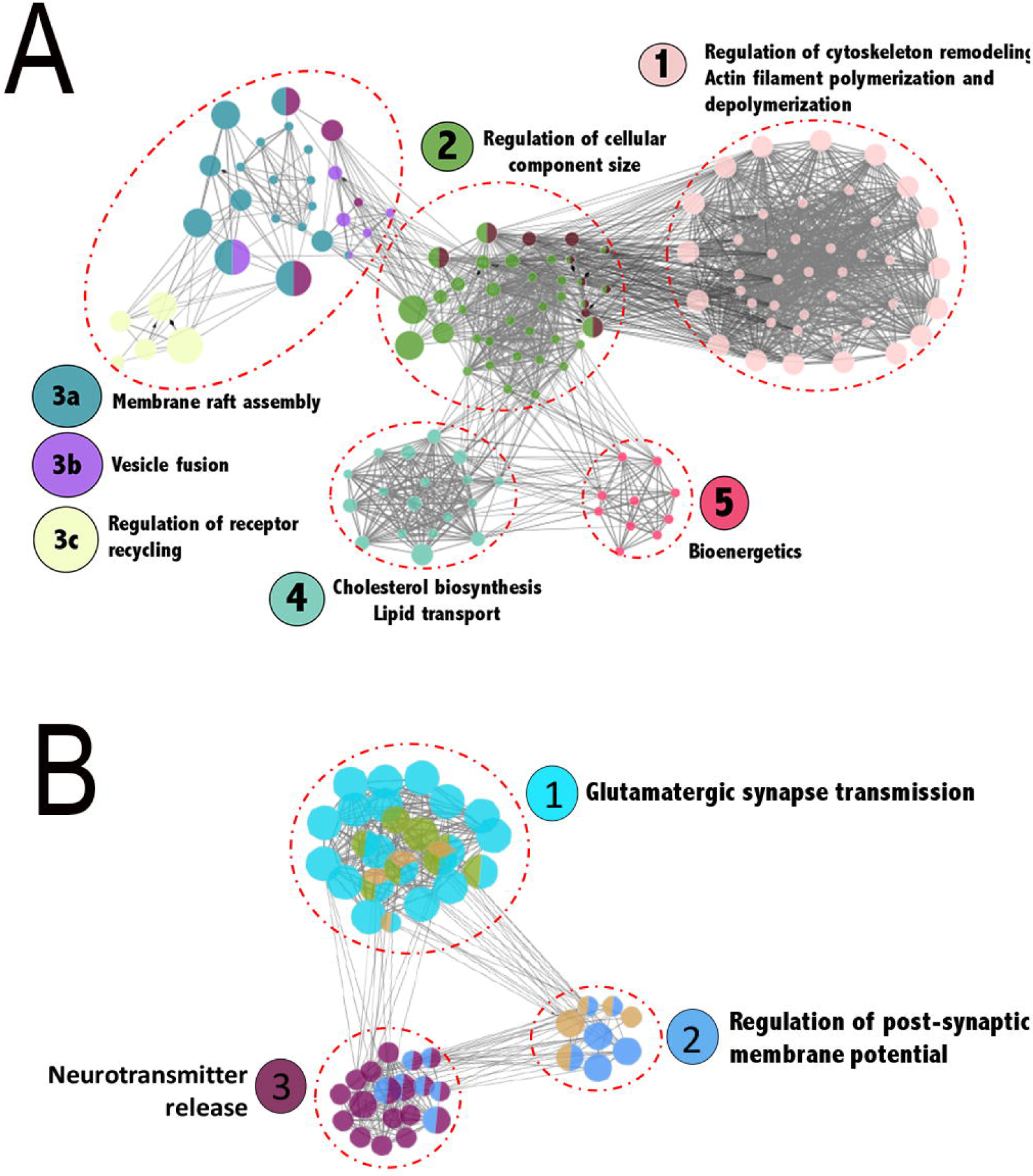
Transcriptional mechanisms underlying KLF6 mediated growth promotion investigated through RNA-Seq analysis. (A) Regulatory network analysis of genes upregulated after KLF6 overexpression revealed sub-networks enriched for distinct functional categories highly relevant to axon growth. Nodes correspond to target genes and edges to multiple modes of interaction (physical, shared upstream regulators, shared signaling pathways and inter-regulation). Node color represents their corresponding functions as denoted in the legends and node size is based on significance for enrichment in functional category. Only significantly enriched GO categories were included in the network analysis (p<0.05, Right-sided hypergeometric test with Bonferroni correction, Cytoscape).

### Functional Synergy between KLF6 and STAT3

As an alternative to clarifying downstream cellular effects, analysis of differential gene expression can also shed light on KLF6 function by identifying additional transcription factors that may functionally interact. We therefore performed motif analysis of the promoter regions of genes upregulated by KLF6 expression, in order to identify factors that could potentially synergize by co-occupying promoters of KLF6-responsive genes. Consistent with regulation by KLF6, the motif most enriched in these promoter regions was the recognition sequence for KLF7/KLF6 itself. Also over-represented were recognition motifs for additional factors including KLF4, SP1, CTCF, and STAT3 (**Fig. 5A**). To test potential functional interactions, we overexpressed these factors singly or in combination with KLF6 in assays of neurite outgrowth. Co-expression of KLF6 with KLF4 has been previously tested (Moore et al., 2009), and we therefore focused here on SP1, CTCF, and STAT3. EGFP reporter was co-expressed with the factors to identify transfected cells, using 1:4:4 plasmid ratios that resulted in approximately 80% co-expression of both test plasmids in EGFP+ neurons (**Fig. 5B**). As expected, single overexpression of KLF6 significantly increased neurite lengths (123.9% of control, p<.01, ANOVA with post-hoc Sidak’s) (**Fig. 5C**). SP1 and CTCF had no effect on neurite outgrowth when expressed singly, and when combined with KLF6 did not differ significantly from the effect of KLF6 alone (p>.05, ANOVA with post-hoc Sidak’s). To test STAT3 function we used two constitutively active forms, based on prior work indicating that active STAT3, but not wildtype, affects neurite growth in this culture system and others (Smith et al., 2011; Mehta et al., 2016). Consistent with prior reports, single overexpression of VP16-STAT3 significantly (albeit modestly) increased neurite lengths to 115.6±2.7% of control, whereas caSTAT3 had no significant effect (106.9±2.8%, p<.05 ANOVA with post-hoc Sidak’s). Remarkably, co-expression of caSTAT3 and VP16-caSTAT3 with KLF6 increased neurite lengths to 161.2.6±5.6% and 175.4±9.6% of control, respectively (p<.01 compared to KLF6 alone, ANOVA with post-hoc Sidak’s) (**Fig. 5B,C**). This degree of neurite outgrowth is well above that predicted by the additive effects of cSTAT3 and KLF6, indicating functional synergy.

**Figure 5.**
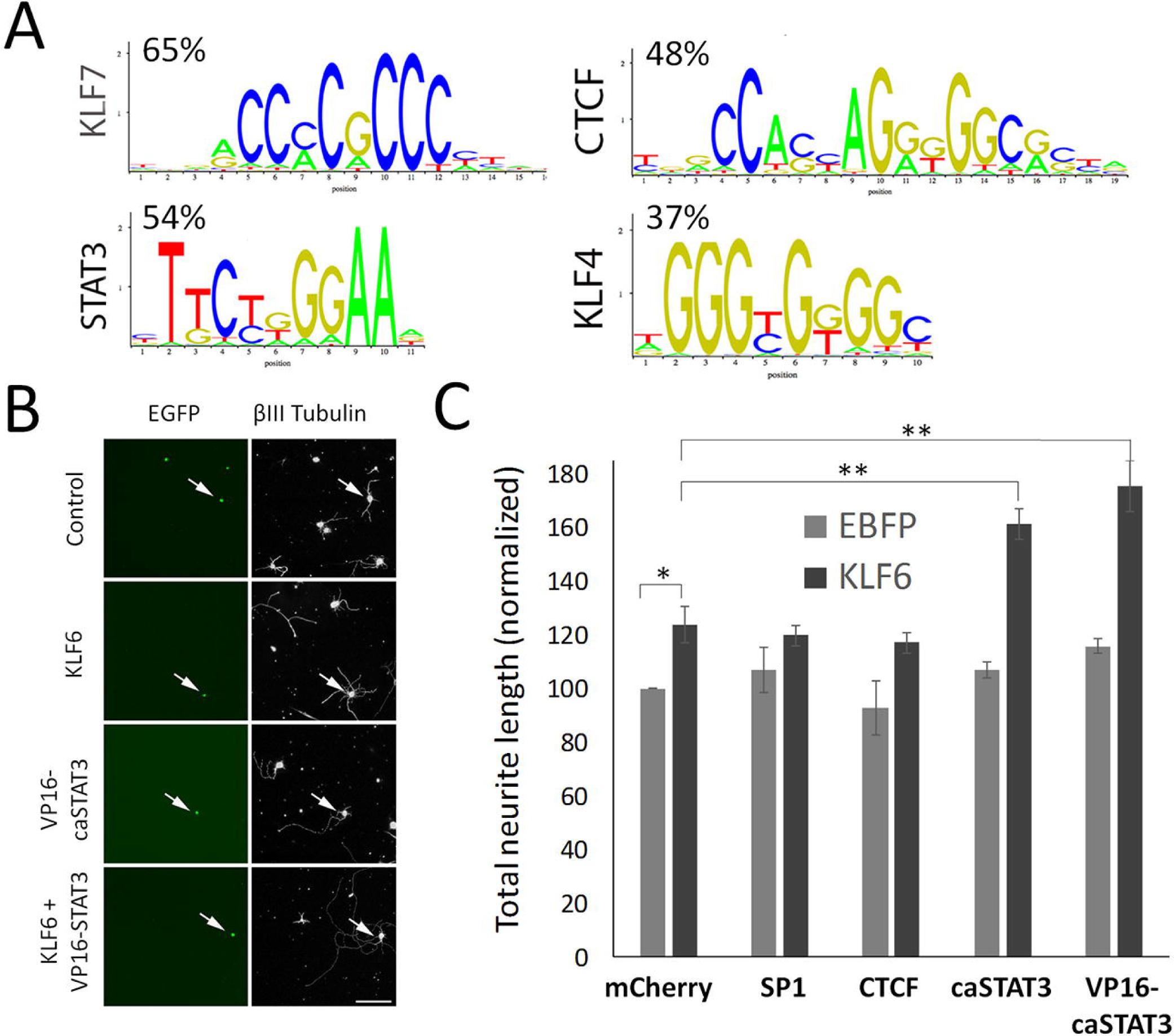
Promoter analysis and phenotypic testing reveals functional KLF6/STAT3 synergy for neurite outgrowth. (A) Transcription factor binding site analysis was performed on the promoters of the 100 genes most upregulated by KLF6 overexpression. Motifs for KLF7 (highly similar to related KLF6), STAT3, KLF4, and CTCF were the most frequently detected motifs (p<.001, Right-sided hypergeometric test with Bonferroni correction, oPOSSUM3). Percentages indicate the percent of promoters in which each motif was detected. (B,C). Postnatal cortical neurons were transfected with KLF6, caSTAT3, VP16-STAT3, or controls for equal plasmid load, along with EGFP reporter. After two days in culture, neurite lengths were measured by automated image analysis (Cellomics) in transfected neurons (EGFP+, green, arrows in B). As expected, KLF6 and VP16-caSTAT3 significantly increased neurite lengths (*p<.05, **p<0.001, p<0.0001, ANOVA with Sidak’s multiple comparisons test, N>200 cells from three replicate experiments). Combined expression of KLF6 with either caSTAT3 or with VP16-caSTAT3 increased neurite length above the level of KLF6, caSTAT3, or VP16-caSTAT3 alone, and above the sum of their individual effects, demonstrating synergistic increases in neurite length. Scale bar is 50µm (B).

**Figure 6.**
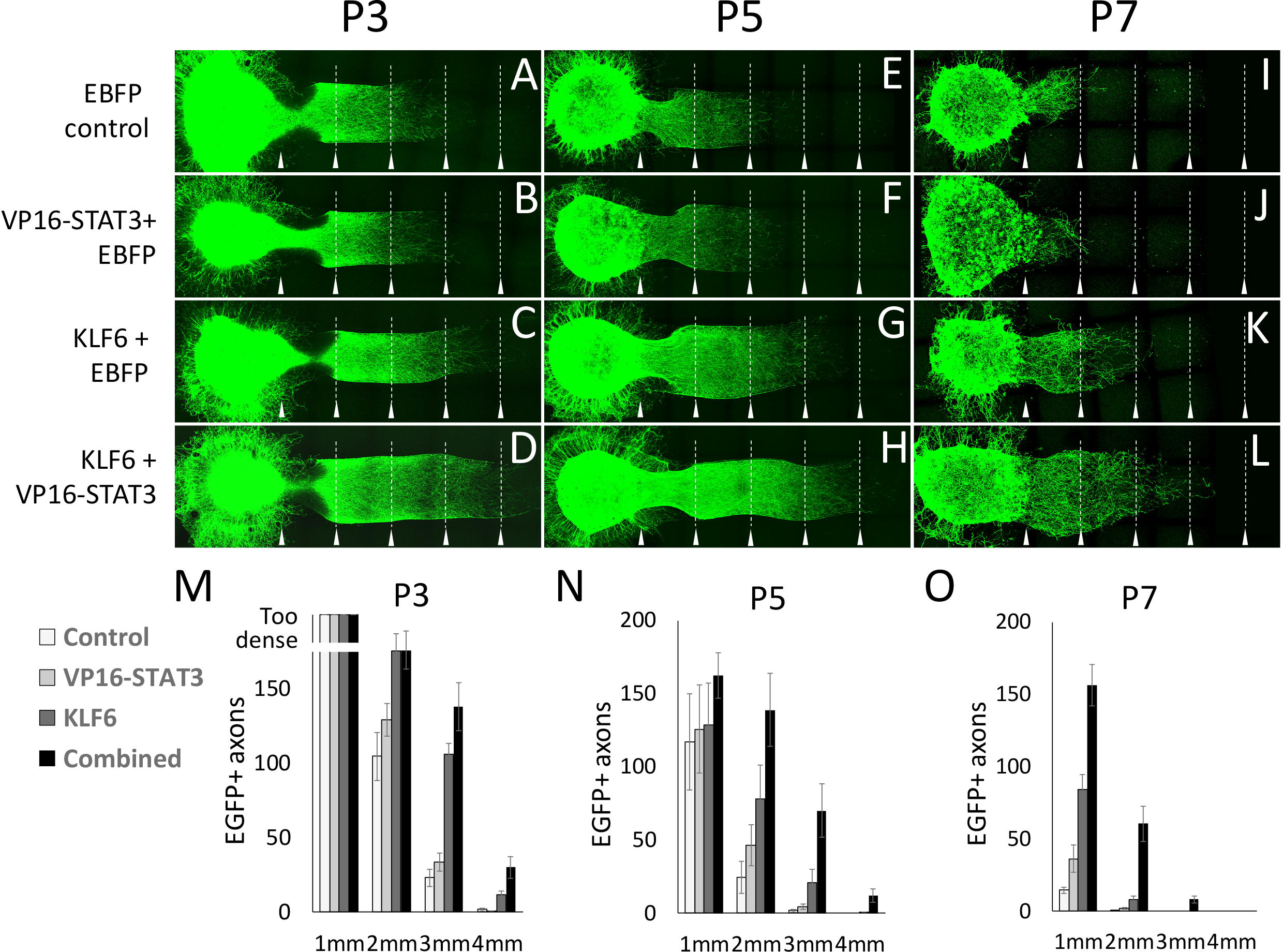
Combined KLF6 and VP16-STAT3 overexpression promotes axon elongation in postnatal cortical neurons. Postnatal cortical cells were dissociated and transfected with an EGFP tracer along with EBFP control, KLF6, VP16-STAT3, or combined KLF6/VP16-STAT3. Transfected cells were reaggregated in matrigel drops, then placed adjacent to a linear column of matrigel that contained no cells. One week later, EGFP+ axons extended as far as 4mm from the edge of the aggregated cells. Baseline axon extension declined in control-transfected neurons between P3 and P7 (compare A, E, I). KLF6 expression increased axon extension, and combined VP16-STAT3/KLF6 expression significantly increased long distance axon growth above KLF6 alone. Arrowheads mark 1mm intervals from the edge of the aggregated cell bodies. N≥6 explants per treatment from two independent experiments.

**Figure 7.**
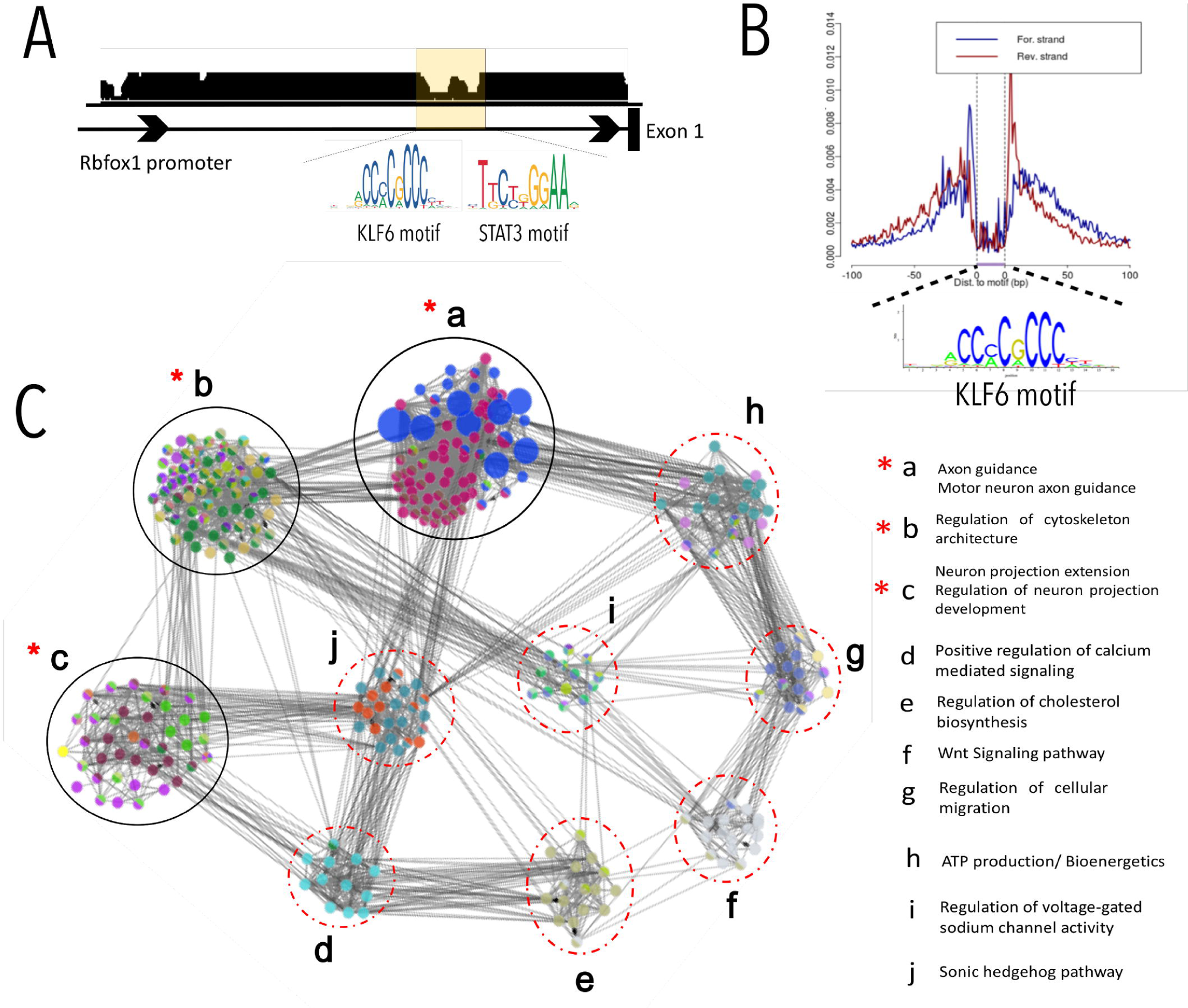
TF footprinting confirmed genome-wide co-occupancy of KLF6/STAT3 in regulatory regions relevant to axon growth. ENCODE consortia ATAC-Seq datasets generated from PO mouse cortex were used for genome-wide TF footprinting to identify regulatory regions bound *in vivo* by KLF6 and STAT3. (A) Representative UCSC browser image of gene promoter region with clear notches within peaks of accessibility that correspond to KLF6 and STAT3 motifs confirming co-occupancy. (B) Aggregate ATAC-Seq footprint for KLF6/7 (shown motif) generated over binding sites within the genome. (C) Regulatory network analysis of genes with deep KLF6/7 footprints revealed 10 sub-networks (a-i) enriched for distinct functional categories highly relevant to axon growth (see legend). Genes showing co-occupancy by both KLF6 and STAT3 were functionally clustered in sub-networks a,b,c (indicated by *-red asterisk) Nodes correspond to target genes and edges to multiple modes of interaction (physical, shared upstream regulators, shared signaling pathways and inter-regulation). Node color represents their corresponding functions as denoted in the legends and node size is based on significance for enrichment in functional category. Only significantly enriched GO categories were included in the network analysis (p<0.05, enrichment – right-sided hypergeometric test with Bonferroni correction).

As a further test of synergy between KLF6 and VP16-STAT3, we tested co-expression in a second growth assay. Postnatal cortical neurons dissociated and transfected as previously, but then re-aggregated in drops of matrigel to create artificial explants. These explants were placed adjacent to lanes of matrigel that contained no cells. The key advantage of this assay is that unlike individually plated cells, aggregated neurons display sustained, long distance axon growth that extends several millimeters. This provides a better model of long distance axon regeneration *in vivo* (**Fig.6A-L**). The assay detected an age-dependent decline in axon extension, with 104.7±16 axons reaching 2mm from P3 explants, 24.4.7±11 from P5, and 0.1±0.1 from P7 (p<.01, ANOVA with post-hoc Sidak’s) (**Fig.6 M-O**). Forced expression of KLF6 increased axon growth at all ages: 175±11.4, 78.1±23.3, and 8.0±2.5 axons reached 2mm from P3, P5, and P7 explants (**Fig.6 M-O**). In contrast, VP16-STAT3 treated neurons showed no increased axon extension in this assay. When VP16-STAT3 and KLF6 were combined they again produced a synergistic increase in axon extension in P5 and P7 explants (138.8±25.0 and and 60.73±12.14 axons reached 2mm) (**Fig.6 N,O**). The combined KLF6/VP16-STAT3 was the only condition in which P7 axons ever extended as far as 3mm (**Fig.6 O**). Thus combined expression of VP16-STAT3 and KLF6 produces synergistic enhancement of axon elongation that partially blocks the age-dependent reduction in axon extension in postnatal cortical neurons.

### Co-occupancy by KLF6 and STAT3 in axon growth gene networks

Transcription factors can functionally synergize through a variety of mechanisms, including physical binding, transcriptional activation of upstream regulators, and co-occupancy of regulatory DNA (Venkatesh and Blackmore, 2017). We considered physical binding, which has been shown between STAT3 and another KLF family member, KLF4 (Qin et al., 2013). KLF6 lacks the STAT3 binding region identified in KLF4, however, and no KLF6/STAT3 interactions are reported in available interaction databases. Regarding cross-activation, recent profiling in oligodendrocytes indicated that KLF6 activity may influence STAT3 by increasing expression of upstream regulators including gp-130 and JAKs (Laitman et al., 2016). Examination of our RNA-seq analysis did not detect upregulation of these genes by KLF6, arguing against a similar mechanism in neurons.

Without strong support for physical binding or transcriptional cross-regulation in our system, we therefore focused on potential co-occupancy of DNA, which was indicated by the prior motif analysis. We performed footprinting analysis of ATAC-seq data from early postnatal cortex, which enables genome-wide identification of loci that are stably bound by TFs (Pique-Regi et al., 2011; Sung et al., 2016) (Fig.11 A,B). This analysis found ~700 deep footprints that significantly correspond to the consensus binding sequence for KLF6/7 in promoter regions or in intragenic enhancer regions (see methods for a discussion of enhancer identification) (**Fig.7 A,B**). We performed network analysis of the corresponding genes, which revealed ten subnetworks. Again consistent with the role of KLF6 in axon growth, three prominent sub-networks were enriched for axon-relevant terms including axon guidance, axon projection, and cytoskeletal organization (**Fig.7C**). We next performed footprint analysis to identify sites of STAT3 binding that occurred within the same promoter/enhancer regions as KLF6 footprints, and found that overall approximately 6% of regulatory regions with KLF6 footprints were also co-occupied by STAT3, a rate that is significantly higher than chance (Wang et al., 2013). Most importantly, genes that displayed STAT3 co-occupancy were not spread evenly across the entire KLF6 network, but instead occurred in only the axon-relevant sub-networks (guidance, projection, and cytoskeleton) (**Fig.7C**). This finding strongly indicates that STAT3 and KLF6 influence axon growth by co-occupying regulatory DNA in growth-relevant gene networks (Venkatesh and Blackmore, 2017).

## DISCUSSION

Transcription factors are emerging as powerful tools to enhance axon regeneration, but likely must be supplied in appropriate combinations for maximal effect. Efforts to identify optimal combinations are hampered by limited understanding of the transcriptional mechanisms that underlie TF-triggered growth. Here we identified KLF6 as an effective pro-regenerative TF, and then integrated transcriptional profiling with recent advances in gene network analyses to identify KLF6-responsive gene modules. We then leveraged these insights to identify STAT3 as a TF that functionally synergizes with KLF6 in promoting axon extension and identified co-occupancy of the two factors in regulatory DNA associated with specific pro-regenerative gene networks. These results offer new insight into transcriptional networks that underlie axon extension and suggest combined KLF6/STAT3 activity as a first step in their reconstruction.

### Network analysis reveals pro-regenerative modules that respond to KLF6

KLFs, a 17-member family of zinc finger transcription factors, have emerged as important regulators of the neuron-intrinsic capacity for axon growth (Veldman et al., 2007; Moore et al., 2009). We previously demonstrated that KLF7 increases corticospinal regeneration *in vivo* when delivered as a transcriptionally active and protein-stabilized mutant (VP16-KLF7), but is ineffective in its wildtype form (Blackmore et al., 2012). Here we show that KLF6, like KLF7, is developmentally downregulated in cortical neurons, and that forced re-expression enhances CST regeneration after spinal injury. Unlike KLF7, however, wildtype KLF6 protein is readily detectable after viral overexpression, indicating that KLF6 protein is not subject to the same regulation as KLF7. Importantly, KLF6 and −7 very likely act through similar transcriptional targets. They share nearly identical DNA-binding domains, indicating similar genomic targeting, and functionally compensate for one another in promoting axon growth (Veldman et al., 2007). Indeed, we showed previously that forced co-expression of KLF7 and KLF6 produces no gain in axon growth above either alone, again indicating a shared mechanism (Moore et al., 2009). Thus, because KLF6’s effects match KLF7’s but do not depend on an artificial VP16 transcription domain, it is likely a better candidate for eventual translation and is a preferable tool to clarify the endogenous transcriptional mechanisms of pro-regenerative TFs.

In recent years, transcriptome profiling approaches have revealed large numbers of genes associated with various forms of endogenous or TF-stimulated axon growth (Li et al., 2015; Siegel et al., 2015; Chandran et al., 2016; Norsworthy et al., 2017). Extracting biological insights from lists of differentially expressed genes, however, remains a non-trivial challenge. One standard approach is to test for overall enrichment of ontological terms, which can provide a first-pass indication of the cellular functions that are impacted by the gene set. One the other hand, this approach leaves unanswered the question of how the gene products might interact, and risks missing functions that exist in subnetworks but which are diluted by whole-set analysis. Accordingly, here we applied a network-based approach in which genes were first clustered into sub-networks according to mutual interactions. This analysis revealed that KLF6-responsive genes broke into five prominent sub-networks. Interestingly, when ontological enrichment analysis was applied to each subnetwork, all five showed significant enrichment for functions that play important but complementary roles in axon growth (Kamiguchi, 2006; Liu et al., 2011; Stuermer, 2011; Tedeschi, 2012; Gomez and Letourneau, 2014; Gordon-Weeks and Fournier, 2014; Ma and Willis, 2015). Similarly, network analysis of genes downregulated upon KLF6 over-expression revealed subnetworks that were highly enriched for synaptic plasticity and neurotransmission. This is consistent with the growing body of evidence that the developmental decline in axon growth capacity is due in part to a neuronal transition from axon extension to synaptogenesis (Tedeschi et al., 2016; Norsworthy et al., 2017; Tedeschi and Bradke, 2017). These results highlight the ability of transcriptional profiling and network analysis to yield new insight into the complementary cellular networks activated by pro-regenerative treatments.

### Combinatorial TF expression for axon growth

It is increasingly appreciated that manipulation of multiple TFs will likely be necessary to evoke a full regenerative response, and recent efforts have sought to identify optimal combinations (Lerch et al., 2014; Fagoe et al., 2015; Chandran et al., 2016; Venkatesh and Blackmore, 2017). TFs can interact through a variety of mechanisms, and the details of these interactions determine the optimal strategies for detecting them (Venkatesh and Blackmore, 2017). For example, prior efforts to identify regenerative TF combinations have focused on direct protein-protein interactions in order to build networks and identify hub TFs (Chandran et al., 2016). However, TFs commonly synergize by targeting overlapping sets of regulatory DNA, without direct physical TF-TF interaction (Wang et al., 2013). Thus a molecular signature of functionally synergizing factors is frequent co-occupancy of promoter regions. Detection of co-occupancy has been leveraged to clarify regulatory TF networks in other biological processes, but has not previously been examined for axon growth (Kidder et al., 2008; He et al., 2011; Wang et al., 2012; Foley and Sidow, 2013; Lee and Zhou, 2013; Liu et al., 2015).

Therefore, starting with the KLF6-responsive gene modules, we analyzed upstream regulatory DNA to identify relevant TFs that may synergize with KLF6 to drive growth. We observed that the recognition motif for STAT3 is highly over-represented in the promoter region of genes that are upregulated by KLF6 overexpression, suggesting a potential interaction between the two factors. STAT3 has been previously shown to have pro-regenerative effects in retinal ganglion cells and CST neurons (Lang et al., 2013; Pernet et al., 2013; Luo et al., 2016; Mehta et al., 2016). Interestingly, transcriptionally active VP16-STAT3, but not wildtype, was recently shown to promote axon growth in postnatal cortical neurons. Consistent with this, we found that VP16-STAT3 produced a modest increase in axon growth in one of the two assays used here. When combined with KLF6, however, VP16-STAT3 produced a dramatic increase in neurite extension in two growth assays, including an explant-based assay of long distance axon growth. Thus, as predicted by bioinformatics modeling, KLF6 and STAT3 synergize to enhance axon growth in developing cortical neurons.

Finally, we took advantage of recent advances in genome-wide digital footprinting to substantiate the prediction that KLF6 and STAT3 co-occupy regulatory DNA of pro-regenerative genes. To our knowledge, this analysis created the first comprehensive in vivo binding map for pro-regenerative TFs during periods of robust axon extension. In line with the RNA-seq based network analysis, we found that KLF6-footprinted genes separated into several sub-networks, some of which were highly enriched in functions relevant to axon growth including neuron projection extension, motor neuron axon guidance, and regulation of neuron projection extension. Importantly, instances of co-occupancy by STAT3 were not distributed randomly across the entire network, but instead occurred exclusively in the growth-relevant subnetworks. This finding lends strong support to the model that KLF6 and STAT3 synergistically enhance axon growth through a mechanism of co-occupancy of regulatory DNA. More generally, this analysis highlights the utility of a comprehensive systems approach to identify cooperative TF regulators of axon growth. Starting from genome-wide expression datasets, integrated network and footprinting analyses offer enormous potential to further elucidate TF regulatory circuits that underlie axon growth.

## TABLE LEGENDS

**Sup_Table 1**. List of differentially expressed genes

**Sup_Table 2**. Quantitative PCR primer sequences.

**Figure S1. KLF6 expression in adult corticospinal tract neurons is insensitive to cervical axotomy**. Mice ten weeks of age received C4/5 dorsal hemisections and injection of the retrograde tracer CTB-488. (A) CTB-488 label (green) in cortex identifies corticospinal tract projection neurons. (B-H) Immunohistochemistry for KLF6 (red) was performed, and the average intensity of KLF6 in corticospinal tract neurons (CTB-488, arrows) was normalized to the average intensity of cells in adjacent, uninjured layers of cortex. KLF6 intensity was dim prior to injury, and no change was detected at any post-injury time-point, indicating an unchanged level of KLF6 expression in injured corticospinal tract neurons (P>.05, ANOVA with Tukey, N=200 cells from 3 animals at each time point). Scale bars are 1mm (A) and 20µm (C-H).

**Figure S2. KLF6 expression correlates with and is sufficient to promote neurite outgrowth *in vitro***. (A,B) Cortical cells were prepared from P7 cortex, maintained in culture for two days, and immunohistochemistry for KLF6 (red) and neuron-specific βIII tubulin (white) performed. More than 10,000 cells were analyzed with a high content screening microscope. A naturally occurring distribution of KLF6 intensities was evident (histogram, A). Neurite outgrowth was averaged for the four quartiles of KLF6 expression. The top two quartiles were significantly longer than the bottom two (p<.01, ANOVA with TUKEY). (C-H). Postnatal cortical neurons were transfected with plasmid encoding EBFP control or KLF6. Overexpression of KLF6 (green) was confirmed by immunohistochemistry (arrows, E), and automated microscopy identified and traced neurites (F). Quantification of more than 10,000 individual neurons confirmed a population-level shift in KLF6 expression after transfection (histogram, G). Overexpression of KLF6 significantly increased average neurite lengths in P7 cortical neurons (*p<.05, ANOVA with Sidak’s multiple comparisons, N>200 neurons in three separate experiments). Scale bar is 50µm.

**Figure S3. Transcriptional mechanisms underlying KLF6 mediated growth promotion investigated through RNA-Seq analysis**. P5 cortical neurons were virally transduced to overexpress either EBFP control/ KLF6 and cultured on laminin substrates for 3 days before RNA extraction and deep sequencing. (A) RNA-Seq data analysis workflow (B) High mapping quality of sequenced reads confirmed library and sequencing quality (C) Representative IGV browser image of genomic locus corresponding to differentially expressed genes (s100a10). (D) Sequencing read distribution across the gene body for treatment groups were monitored to ensure optimal library and sequencing quality. Differential gene expression analysis identified 454 genes that significantly differ between control and KLF6 groups (p-value<0.05, FDR<0.05). 55% of the transcripts were upregulated and 45% of the transcripts were down-regulated in response to KLF6 over-expression. (E) Unsupervised hierarchical clustering showed gene expression signatures in control/ KLF6 groups and tight clustering among replicates.

**Figure S4. qPCR validation of transcriptional changes predicted by RNAseq**. (A) Shows a schematic of KLF6 protein, indicating lysine residues shown previously to be subject to acetylation. Asterisks indicate lysines mutated to Q or R. (B,C) P7 cortical neurons were transfected with EGFP reporter and control mCherry, wildtype KLF6, or KLF6 with lysines 209 and 213 mutated to glutamine (Q) or arginine (R). After two days in culture, high content microscopy quantified neurite length in transfected neurons, and qPCR quantified expression of putative KLF6-regulated genes. (B) Wildtype KLF6 and KLF6(K,R), but not KLF6(K,Q) significantly increased neurite length (****p<0.0001, ANOVA with Tukey’s multiple comparisons, N=>200 cells from three replicate experiments). (C) Wildtype KLF6 and KLF6(K,R) increased the expression of predicted KLF6-regulated genes, while KLF6(K,Q) did not (N=3, ****p<0.0001, **p<0.001, RM 2-way ANOVA with Sidak’s multiple comparisons test).

## Acknowledgments

This work was supported by grants from NINDS, the International Spinal Research Trust, The Bryon Riesch Paralysis Foundation. The authors acknowledge ENCODE consortia and specifically The Bing Ren lab at UCSD for generating the ATACC-seq datasets used in this study. The authors declare no competing financial interests.

